# The future abundance of key bird species for pathogen transmission in the Netherlands

**DOI:** 10.1101/2024.10.14.617534

**Authors:** Martha Dellar, Henk Sierdsema, Peter M. van Bodegom, Maarten Schrama, Gertjan Geerling

**Affiliations:** Deltares, Daltonlaan 600, 3584BK, Utrecht, The Netherlands; Institute of Environmental Sciences, University of Leiden, Van Steenis Building, Einsteinweg 2, 2333CC, Leiden, The Netherlands; Sovon Dutch Centre for Field Ornithology, Toernooiveld 1, 6525 ED, Nijmegen, The Netherlands

## Abstract

Wild birds serve as reservoirs and vectors for many different pathogens and changes in their distribution and abundance due to environmental change will influence disease risk. We study three species which are highly abundant in north-western Europe and which can transmit a wide range of diseases including avian influenza and West Nile virus: blackbirds, mallards and house sparrows. Using the Netherlands as a case study, we created random forest models for predicting the distribution and abundance of these species, both now and in the future. Climate, land use and vegetative cover were all important predictors of bird abundance. The three species had different spatial distributions, largely related to their preferred habitat and food availability. In the future, mallard and house sparrow populations were predicted to increase, while there was little change for blackbirds. Quantifying the consequences of these abundance changes is complicated as there are many factors to consider, however increased pathogen reservoirs will likely increase disease risk and changes in distribution may affect local outbreak risk. The future abundance maps created in this study, and the methods used to create them, will be useful tools for disease modellers and policymakers to estimate future disease risk and to plan accordingly.

## Introduction

Wild birds can serve as both reservoirs and mechanical vectors for many different pathogens, with health implications for humans, domestic animals, other wild animals and the birds themselves (Levison, 2016; Reed et al., 2003; Tsiodras et al., 2008). Different bird species transmit different pathogens, and for many of these pathogens it is not known which species play the most important role in their transmission. In northwestern Europe, recent outbreaks of Usutu virus (USUV), West Nile virus (WNV) and avian influenza (AI) have highlighted the risk of bird-borne pathogens and the ease with which they can spread (Caliendo et al., 2024; Oude Munnink et al., 2020; Sikkema et al., 2020; Vlaskamp et al., 2020).

Key to a better understanding of the risks of bird-borne pathogens is an understanding of the distribution and abundance of key bird species. The recent Bioscore2 project made predictive abundance maps for a suite of different bird species across Europe (Hendriks et al., 2016). These and other studies have shown that changes in both climate and land use lead to changes in the distribution and abundance of birds (Howard et al., 2015; Jetz et al., 2007; Kampichler et al., 2012). This implies that future risks of bird-borne diseases may strongly deviate from current spatial patterns in risk. However, these studies did not focus on key bird species in relation to diseases and have generally not considered knock-on effects on pathogen transmission. Given the lack of knowledge on the future distribution of key bird species for pathogen transmission, it is highly valuable to have future predictions for the distribution and abundance of such species. Such predictions can be used in modelling studies to estimate future disease risk (De Wit et al., 2024) and provide tools for acting upon such predicted changes for risk control.

We know several species which are likely to make a significant contribution to the spread of certain diseases in northwestern Europe; these are common blackbirds (*Turdus merula*), house sparrows (*Passer domesticus*) and mallards (*Anas platyrhynchos*) (Ashraf et al., 2015; Beer et al., 2021). By focusing on a particular region and a few key species, we can gain a better understanding of the role of shifting bird distributions on pathogen transmission. These species all have large breeding populations in northwestern Europe, are collectively capable of transmitting a wide range of pathogens and were not included in the Bioscore2 project. They are also all commonly found in urban and peri-urban areas, meaning that they will often be in close contact with people, with possible consequences for human health. Blackbirds are part of the *Turdidae* family. They are highly susceptible to USUV, which has caused significant reductions in the European population in recent years (Siljic et al., 2023; van den Bremer & van Turnhout, 2021). Blackbirds are also known to be susceptible to avian malaria (Agliani et al., 2023), competent hosts of WNV (Angelou et al., 2021) and a key reservoir species for Lyme borreliosis in central Europe (Taragel’ová et al., 2008). House sparrows are part of the *Passeridae* family. They are known to be competent hosts for WNV, AI and avian malaria (Medeiros et al., 2016; Shriner & Root, 2020), as well as bacterial diseases such as *Escherischa coli* and *Salmonella spp.* (Vilela et al., 2012). Mallards are part of the *Anatidae* family. They are known to be competent hosts for WNV and AI and they have the potential to spread pathogens over long distances as part of their migratory movements (Kilpatrick et al., 2007; Pérez-Ramírez et al., 2014; van Toor et al., 2018).

In this study, we aim to determine how climate change and land use change affect the distribution and abundance of these three key bird species. We will also consider the effects these changes might have on pathogen transmission. We use the Netherlands as a case study, since there are both large amounts of bird data available and extensive environmental data. In addition, new detailed socio-economic scenarios have recently been published for this country (Dellar et al., n.d., 2024), providing detailed information on potential future land use changes. For all three species under consideration, it is estimated that there are at least 180,000 breeding pairs in the Netherlands, and for house sparrows there may be up to a million (Sovon, 2023). Also, the Netherlands has recently experienced a significant USUV outbreak as well as its first introduction of WNV (Sikkema et al., 2020; van den Bremer & van Turnhout, 2021). However, all these species are common throughout northwestern Europe and the results from this study should be widely applicable. As well as providing detailed future predictions for the Netherlands on the abundance and distribution of these species, our results will paint a general picture on how future pathogen transmission may be affected by climate- and land use-induced changes in bird abundance.

## Methods

### Overview

We trained a random forest model and used this to make predictions for the distribution and relative abundance of blackbirds, mallards and house sparrows in the Netherlands, for both the current situation and 2050, based on different future scenarios. To train the model, we used bird data from both the Netherlands and France, since the current French climate is similar to the future Dutch climate.

### Bird abundance data

Bird data for the three species under consideration for the Netherlands was taken from the Meetnet Urbane Soorten (MUS), Meetnet Agrarische Soorten (MAS) and Broedvogel Monitoring Project (Common Bird Census, CBC) datasets (Schoppers et al., 2020; Teunissen et al., 2019; Vergeer et al., 2023). For MUS and MAS, point count data is collected during three-four visits in the breeding season by volunteers in urban and rural areas respectively. They count all the breeding birds they see during a five- (MUS) or ten- (MAS) minute period. For MUS, measurements are taken between one hour before and two hours after sunrise, while for MAS they are taken up to five hours after sunrise. Some measurements were taken outside the breeding season and these were excluded. For CBC, volunteers walk a set route five to ten times within a prescribed area, recording all the birds, or a preset selection of species, they see or hear. This only happens during the breeding season and generally during the first five hours after sunrise. The data is then converted to a 250m grid. The monitoring areas cover all major land use types in the Netherlands. The breeding seasons of the three species used are shown in table 1. We excluded measurements taken in the same location in the same year within each dataset, keeping just the maximum value.

**Table 1:** Breeding season per species (van Turnhout, 2011) Species Breeding season.

| Species | Breeding season |
| --- | --- |
| Blackbird | March 1<sup>st</sup> – July 15<sup>th</sup> |
| Mallard | April 1<sup>st</sup> – June 10<sup>th</sup> |
| House sparrow | April 10<sup>th</sup> – June 19<sup>th</sup> |

Bird data for France was taken from the Suivi Temporel des Oiseaux Communs (STOC) dataset (Vigie-Nature, 2021). Point count data is collected by volunteers during the breeding season, counting all the birds they see or hear during a five-minute period. Measurements are taken one to 4 hours after sunrise. Again, we excluded measurements taken in the same location in the same year.

We used data from France since the current French climate is comparable to the Dutch climate in 2050 (Bastin et al., 2019). We only took French bird data from areas where the climate was sufficiently similar. To estimate the future Dutch climate, we used summarised statistics from future climate scenarios produced by the Dutch Meteorological Association (KNMI, 2023). These are included in the supplementary materials (‘Climatic ranges’) and include average minimum, mean and maximum temperatures as well as total rainfall for different seasons, with a total of ten variables. For each climatic variable, a range was given for 2050. We selected areas of France which fell in or near these ranges, while still capturing an area which covered the more extreme values which are expected due to increased future climatic variability. To this end, we chose the areas of France where, for at least five of the ten variables, the average climate was within the given range, plus or minus an additional margin equal to the range’s length, to account for an increase in extreme values. This meant that we excluded those areas where the climate was dramatically different to the Netherlands. French climatic data was taken from the E-OBS dataset (Cornes et al., 2018), using ensemble mean data on a 0.1° grid. The French area deemed to have a comparably similar climate to the Netherlands in 2050 is shown in figure 1. Finally, we removed any remaining French datapoints with an altitude above 350m, since this is the highest point in the Netherlands and we are only interested in French areas which are comparable.

**Figure 1:**
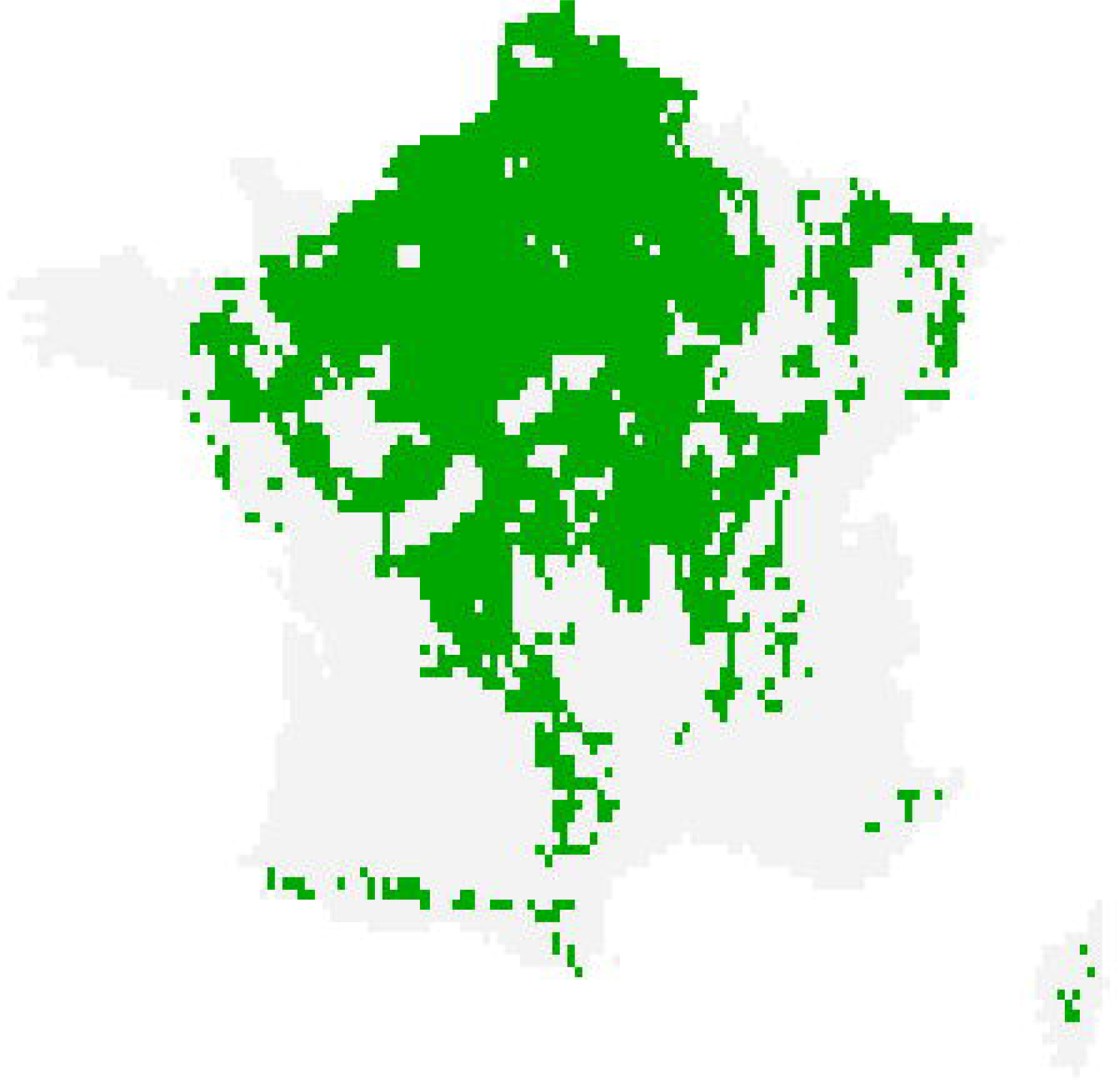
French area from which bird data was taken for our study (green area). The current climate and elevation in this area was deemed sufficiently similar to the Dutch climate and elevation in 2050.

Both the Netherlands and France have experienced recent Usutu outbreaks which have had a particularly large effect on the blackbird population (Lecollinet et al., 2016; Oude Munnink et al., 2020). To avoid this affecting our results, we discarded blackbird data from after 2015 for France, and from after 2016 for the Netherlands. The final number of datapoints per dataset is shown in table 2 and their locations are shown in figure 2.

**Figure 2:**
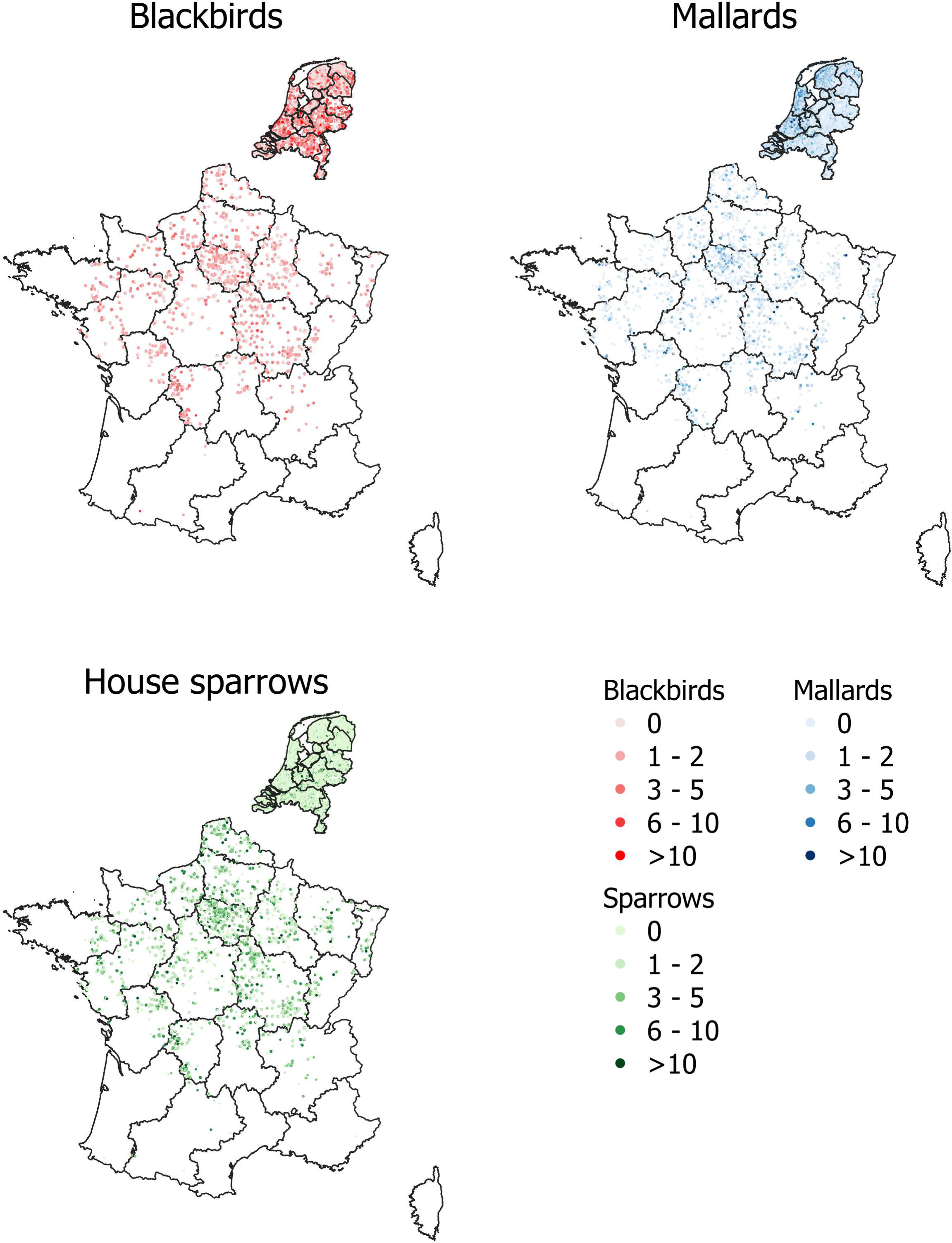
Point locations of observational data used to train our model. Darker shades indicate that more birds were observed.

**Table 2:** Number of datapoints per dataset.

| Dataset | Country | Number of presences | Number of absences | Time period |
| --- | --- | --- | --- | --- |
| Meetnet<br>Urbane Soorten<br>(MUS) | Netherlands | Blackbirds: 50,794<br>Mallards: 14,186<br>House sparrows:<br>24,758 | Blackbirds: 9,123<br>Mallards: 44,476<br>House sparrows: 34,386 | 2007-2023 |
| Meetnet<br>Agrarische<br>Soorten (MAS) | Netherlands | Blackbirds: 1,228<br>Mallards: 2,630<br>House sparrows: 626 | Blackbirds: 3,215<br>Mallards: 1,807<br>House sparrows: 3,813 | 2000-2023 |
| Common Bird<br>Census (CBC) | Netherlands | Blackbirds: 32,742<br>Mallards: 119,431<br>House sparrows:<br>10,798 | Blackbirds: 40,382<br>Mallards: 133,561<br>House sparrows:<br>194,351 | 1983-2023 |
| Suivi Temporel<br>des Oiseaux<br>Communs<br>(STOC) | France | Blackbirds: 33,986<br>Mallards: 2,072<br>House sparrows:<br>11,110 | Blackbirds: 11,653<br>Mallards: 3,522<br>House sparrows: 1,191 | 2001-2023 |

### Covariate data

We used a wide range of covariates for estimating the distribution and abundances of the key bird species, based on literature (Hendriks et al., 2016; Howard et al., 2015; Kampichler et al., 2012) and expert advice from the Dutch Centre for Field Ornithology (Sovon). Climatic data was taken from the E-OBS dataset (Cornes et al., 2018), using ensemble mean data on a 0.1° grid. Climatic parameters were calculated over two time periods: the breeding season for the year the measurement was taken (see table 1) and the winter preceding the breeding season (December 1^st^ – February 28^th^). Parameters used were mean, minimum and maximum temperature, temperature range, number of days with a minimum temperature below 0°C, total precipitation, precipitation coefficient of variation and total precipitation during the driest four-week period. While many other studies have used very general climate data (Hendriks et al., 2016; Howard et al., 2015; Kampichler et al., 2012), often averaged over thirty-year periods, it has been found that using shorter time scales (i.e. weather), as we do here, produces better results when modelling bird distributions (Reside et al., 2010).

Land use data was taken from the CORINE dataset. This is available on a 100m grid for the years 1990, 2000, 2006, 2012 and 2018. For each bird observation, we used land use data from the closest CORINE year. CORINE uses 44 different land use classes. Many of these are quite similar and it is efficient to group them together. The grouping we used is provided in the supplementary materials (‘Deriving future land use’, table S1). Since birds can travel a reasonable distance, different sized buffers around each observation were considered. The proportions of each land-use type within a radius of 500m, 1km and 2km of each observation were used as covariates in our model. We also used tree density and the proportion of grass cover within these same radii. These were taken from the Copernicus Land Monitoring Service and were available on a 100m grid (CLMS, 2020a, 2020b).

We also used the proportions of clay and sand in the soil, from the dataset of topsoil physical properties for Europe on a 500m grid (Ballabio et al., 2016) provided by the European Soil Data Centre (ESDAC, n.d.; Panagos et al., 2012). Altitude was also included and was taken from the European Environment Agency’s elevation map on a 1km grid (EEA, 2016). Finally, we included the dataset from which the training data was taken, to account for variations between our bird data sources.

### Model generation

We used the SDMaps package (Kampichler et al., 2020) in R (R Core Team, 2021), which combines regression-type statistics with spatial interpolation techniques to model species distributions and abundances (Hengl et al., 2009; Sierdsema & van Loon, 2008). We opted to use a random forest model and to produce predictions on a 1km grid.

### Predictions

For each species, we made predictions for the time periods 1991-2020 and 2036-2065. These predictions show relative abundance during the breeding season. It is not possible to calculate absolute abundance from the available data so the results for each species were scaled to have a maximum of 1. Altitude and soil parameters remained unchanged when making future predictions. Land use parameters, grass cover and tree density were derived from Dutch One Health SSPs (Shared Socio-economic Pathways) (Dellar et al., n.d., 2024). See the supplementary materials (‘Deriving future land use’) for full details of how this derivation was performed. The Dutch One Health SSPs are socio-economic scenarios recently developed for the Netherlands with a focus on health. They are based on the global SSP scenarios (O’Neill et al., 2017) and include SSPs 1, 3, 4 and 5. They include land use maps for 2050. Future climate data was taken from scenarios published by the Dutch Meteorological Organisation (KNMI, 2023). They provide six scenarios: low, medium and high emissions, each with a wet and dry variant. The different emissions levels are approximately equivalent to RCPs (Representative Concentration Pathways) 2.6, 4.5 and 8.5 respectively. Each scenario is the result of an 8-model ensemble and uses 1991-2020 as a reference period. The scenarios are on a 12km grid, which we converted to a 1km grid using bilinear interpolation. We paired the low emissions scenario with SSP1, medium emissions with SSP4 and high emissions with SSPs 3 and 5. We averaged over the eight ensembles and also averaged the wet and dry variants.

### Statistical analysis

SDMaps provides a range of different measures for assessing the quality of a model, including mean average error, mean forecast error, root mean-squared error, Pearson correlation and explained variance. It also provides the variable importance (as a percentage) for all covariates. We tested for statistically significant differences between our predictions for different species and scenarios using an ANOVA. We confirmed the assumptions of normality and homogeneity of variance of the residuals and then performed the Bonferroni multiple-comparison test.

### Uncertainty analysis

We calculated the effects of uncertainty in the model generation process. To do this we used a bootstrapping procedure with 200 iterations (trial and error showed this was sufficient to achieve convergence). We made predictions for the 1991-2020 period for each iteration and then calculated the 95% confidence interval per grid square across the iterations.

## Results

### Model performance

The model performance metrics of the random forest models for each of the three species are shown in table 3. All models performed well though the mallard model performed slightly less well than the other two. All models had a slight tendency to over-predict abundance.

**Table 3:** Model performance metrics (MAE: Mean Average Error, MFE: Mean Forecast Error, RMSE: Root Mean-Squared Error)

| Performance metric | Blackbirds | Mallards | House sparrows |
| --- | --- | --- | --- |
| MAE | 0.325 | 0.236 | 0.340 |
| MFE | -0.012 | -0.015 | -0.020 |
| RMSE | 0.544 | 0.603 | 0.896 |
| Pearson correlation | 0.954 | 0.791 | 0.726 |
| Intercept | -0.231 | -0.158 | -0.170 |
| Slope | 1.127 | 1.204 | 1.146 |
| Explained variance | 69.7 | 53.8 | 60.2 |

### Variable importance

We grouped the covariates by type and summed the variable importances within each group, to get a clear picture of what type of variables were the most important. The results are shown in table 4. The full results, including partial dependence plots, are available in the supplementary materials. Land use was always most important. For mallards and house sparrows, climate was the second most important, but for blackbirds it was vegetative cover.

**Table 4:** Grouped variable importance for each bird species model. The numbers in brackets indicate the number of variables in each group.

|  |  | Blackbirds |  | Mallards |  | House sparrows |  |
| --- | --- | --- | --- | --- | --- | --- | --- |
| Climate | Temperature (10) | 20.36 | 12.44 | 35.09 | 21.43 | 24.49 | 15.13 |
|  | Precipitation (6) |  | 7.92 |  | 13.65 |  | 9.36 |
| Land use | Urban (12) | 37.05 | 14.19 | 36.35 | 11.68 | 36.86 | 11.84 |
|  | Agriculture (12) |  | 10.91 |  | 13.83 |  | 14.71 |
|  | Nature (9) |  | 6.93 |  | 5.35 |  | 6.08 |
|  | Water and wetlands (9) |  | 5.02 |  | 5.49 |  | 4.24 |
| Vegetative cover | Grass cover (3) | 22.37 | 8.64 | 15.97 | 7.43 | 18.75 | 8.01 |
|  | Tree cover (3) |  | 13.73 |  | 8.55 |  | 10.74 |
| Other | Soil content (2) | 20.22 | 7.84 | 12.59 | 6.59 | 19.90 | 5.66 |
|  | Altitude (1) |  | 4.03 |  | 4.45 |  | 5.04 |
|  | Dataset ID (4) |  | 8.36 |  | 1.55 |  | 9.20 |

### Current abundance

The relative breeding season abundance for the three species for the period 1991-2020 is shown in figure 3. Blackbirds were generally found to be more abundant in the eastern part of the country though the highest abundances were in urban areas and along the western coast. Mallards on the other hand showed higher abundances in the west. Sparrows were more evenly distributed across the country, though abundances were slightly higher in the south-east, particularly in south-eastern cities, and were noticeably lower in large natural areas.

**Figure 3:**
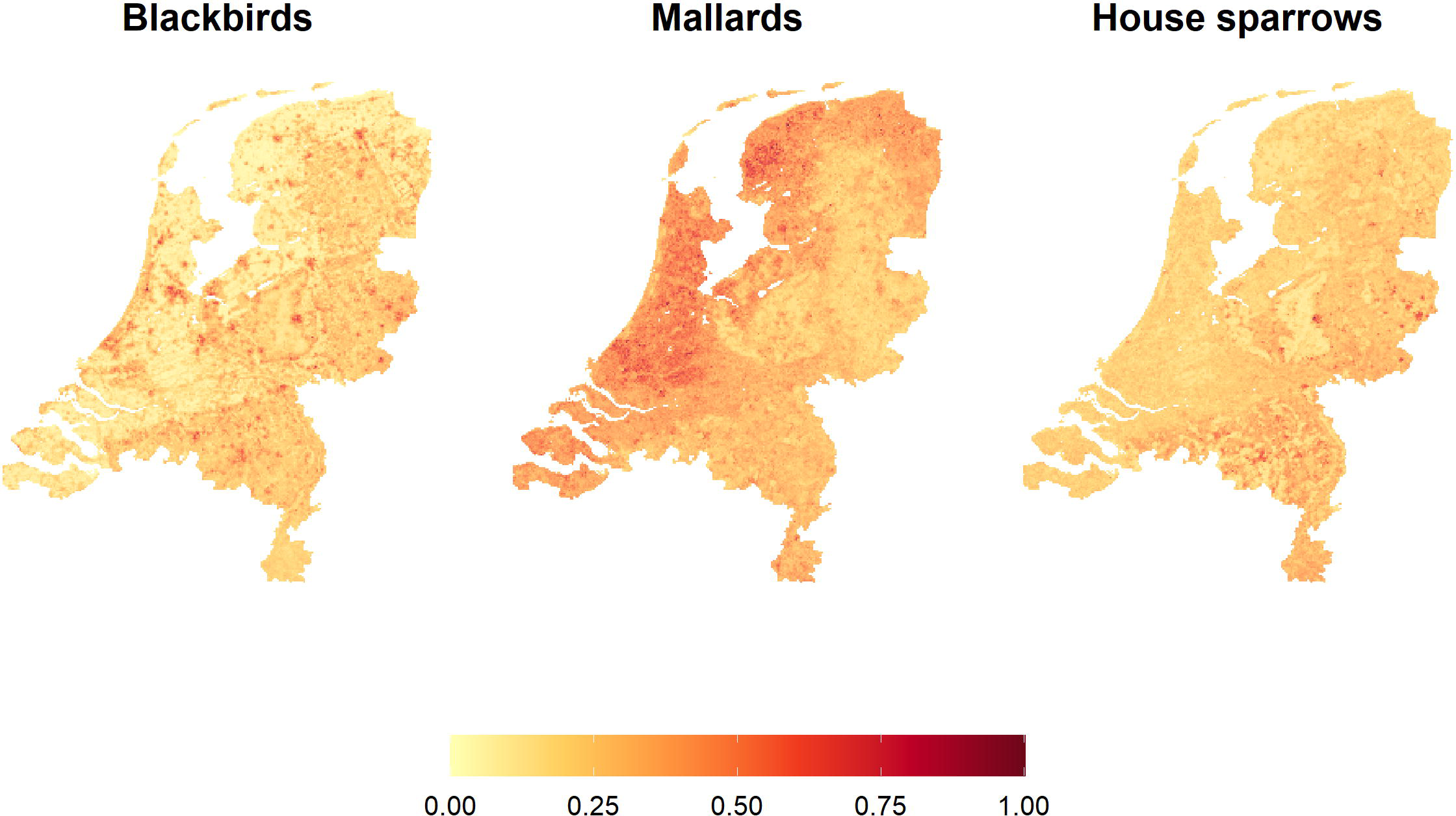
Relative breeding bird abundance during breeding season for the period 1991-2020

### Future abundance

The change in relative abundance between the periods 1991-2020 and 2036-2065 is shown in figure 4 and summarised in table 5. In general, blackbird numbers are expected to stay roughly the same, while the abundances of mallards and house sparrows are expected to increase. Increases are generally larger (relative to current values) in places where abundance is currently low. Future abundance was always significantly higher (p < 0.0001) than in the reference period, except for blackbirds for SSPs 3 and 4, where there was no significant difference. The results for the different scenarios tend to look quite similar, though the differences were mostly statistically significant. Within each species, the differences between scenarios always had p < 0.0001, with the following exceptions: for blackbirds SSP1 vs 5 and SSP3 vs 4 were not significant, for mallards SSP 1 vs 4 and SSP1 vs 5 had significant differences with p < 0.05, and for sparrows SSP3 vs 4 were significantly different with p < 0.001. Within scenarios, the difference between species was always significant with p < 0.0001.

**Figure 4:**
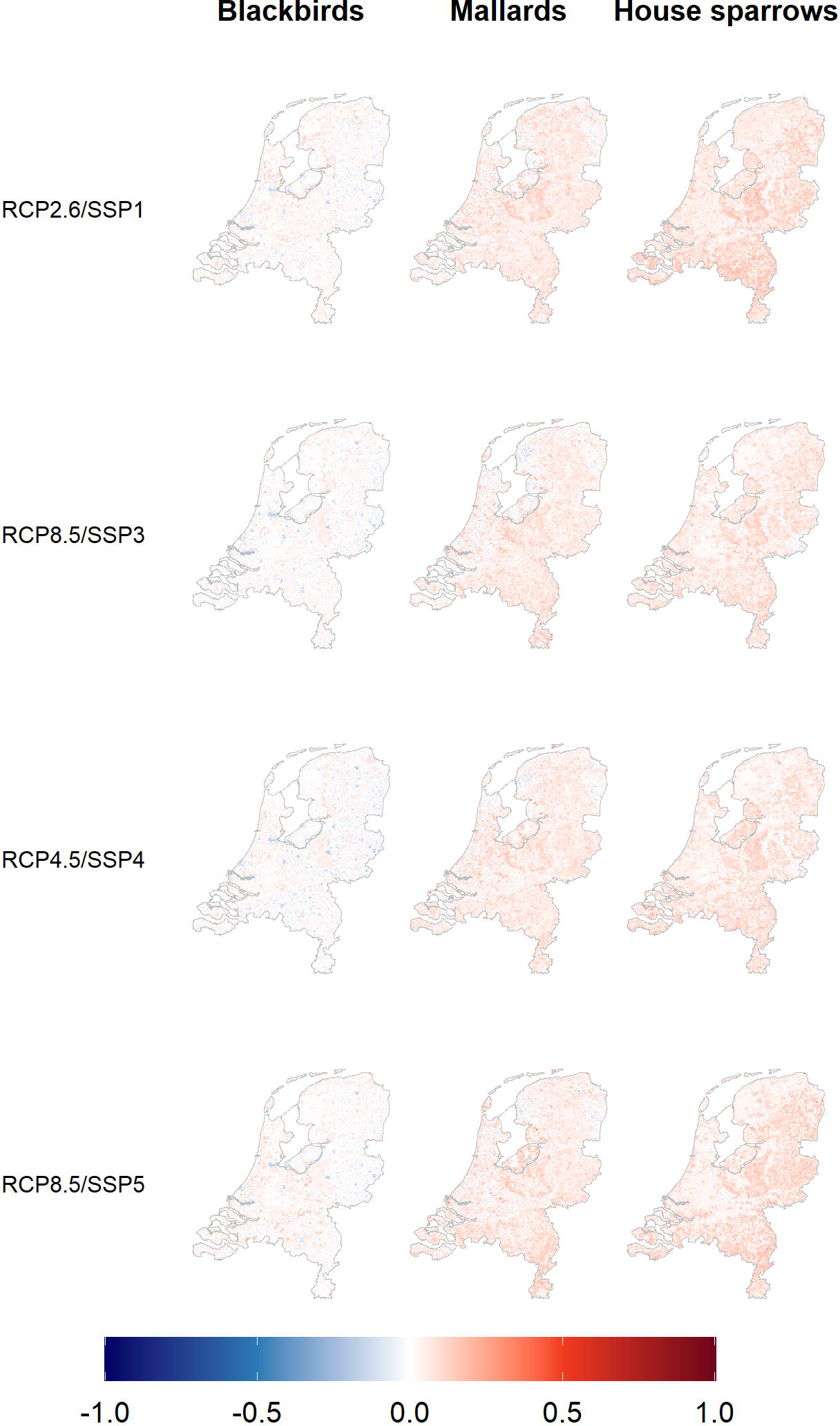
Change in relative abundance of three bird species during the breeding season from present (1991-2020) to future (2036-2065) for each of 4 scenarios. Future distributions were scaled by the same factors as current distributions (Figure 3) and the difference between these and current distributions was calculated. RCP (Representative Concentration Pathway) indicates the climate scenario used and SCP (Shared Socio-economic Pathway) indicates the socio-economic scenario used. White indicates no change from the current situation, while blue indicates a decrease in abundance and red indicates an increase.

**Table 5:**
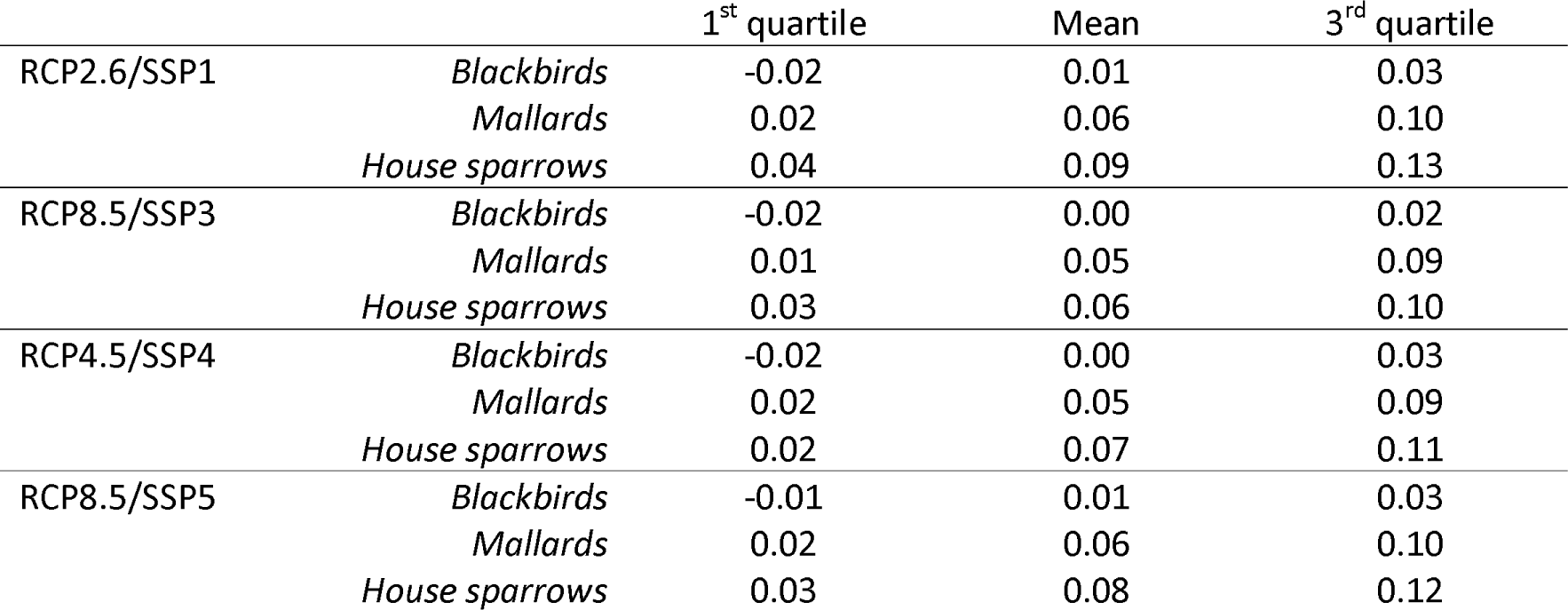
Summary of changes in relative abundance from present (1991-2020) to future (2036-2065). A value of 0 indicates no change. Other values are proportions of the maximum abundance across the country for the present period.

|  |  | 1<sup>st</sup> quartile | Mean | 3<sup>rd</sup> quartile |
| --- | --- | --- | --- | --- |
| RCP2.6/SSP1 | *Blackbirds* | -0.02 | 0.01 | 0.03 |
|  | *Mallards* | 0.02 | 0.06 | 0.10 |
|  | *House sparrows* | 0.04 | 0.09 | 0.13 |
| RCP8.5/SSP3 | *Blackbirds* | -0.02 | 0.00 | 0.02 |
|  | *Mallards* | 0.01 | 0.05 | 0.09 |
|  | *House sparrows* | 0.03 | 0.06 | 0.10 |
| RCP4.5/SSP4 | *Blackbirds* | -0.02 | 0.00 | 0.03 |
|  | *Mallards* | 0.02 | 0.05 | 0.09 |
|  | *House sparrows* | 0.02 | 0.07 | 0.11 |
| RCP8.5/SSP5 | *Blackbirds* | -0.01 | 0.01 | 0.03 |
|  | *Mallards* | 0.02 | 0.06 | 0.10 |
|  | *House sparrows* | 0.03 | 0.08 | 0.12 |

### Model uncertainty

Uncertainty due to the model generation process was calculated. Using the same scaling as for the current abundance maps (figure 3), the average confidence intervals for the whole country were ±0.04, ±0.08 and ±0.06 for blackbirds, mallards and house sparrows respectively. Detailed results are available in the supplementary materials.

## Discussion

We created random forest models to predict the relative abundance of three bird species (blackbirds, mallards and house sparrows) which are likely to be especially relevant for disease transmission in Northwestern Europe. We used these models to predict how the abundance of these species will change under different future scenarios, which included climate, land use and vegetation factors.

Our results show that, besides climate, land use and vegetative cover (and predicted changes therein) are particularly important predictors of bird abundance, with the latter two being more important than climate. Other studies at a European scale have found similar results (Hendriks et al., 2016; Howard et al., 2015; Kampichler et al., 2012), though have often found that climate is the most important predictor. This might be explained by the fact that the Netherlands is a small country and experiences relatively little spatial climatic variation, while at the same time, the high population density (compared with most other countries) leads to a higher prevalence of anthropogenic land use types. Temperature was more important than precipitation in predicting bird abundances, possibly because it has many direct and indirect effects on birds, influencing food availability, breeding success, migration, habitat, predators and competition, among others (Howard et al., 2015). Similarly, land use and vegetative cover have significant effects on both food availability and nesting opportunities (Kampichler et al., 2012). This highlights the need to include a wide range of factors when investigating bird abundance and disease risk.

Looking at current abundance, there is a different pattern for each species. Blackbirds and sparrows have higher abundances in the eastern parts of the country, where there is high soil sand content and greater tree cover, while mallards have higher abundances in the west, where there is high soil clay content and more urban areas. Blackbirds also have very high abundance in urban areas throughout the country, while house sparrows seem to prefer the eastern cities and to particularly avoid large natural areas. These results are in line with previous research. Blackbirds have been becoming increasing urbanised for several decades in the Netherlands, largely as an alternative to migration. This is thought to be due to a combination of climate change and urban areas offering a warmer microclimate and better winter food availability (Evans et al., 2010; Møller et al., 2014; Van Vliet et al., 2009; von dem Bussche et al., 2008). Blackbirds have also been found to prefer forests and woodlands to more open areas as this provides them with a better breeding habitat, which explains their higher abundances in the east than the west (Hatchwell et al., 1996; von dem Bussche et al., 2008). The distribution patterns relating to soil type may be due to indirect effects from relationships between soil and landscape features, or there may be more direct effects. For example, food availability may be a factor, since earthworm density is related to soil type (Rutgers et al., 2016). Other studies looking at mallard distribution have consistently found a strong positive relationship between abundance and the presence of water bodies and wetlands (Barker et al., 2014; Herbert et al., 2018; Milsom et al., 2000; Newbold & Eadie, 2004). While we found a positive relationship with all water and wetland types, this was not a particularly influential factor in our model. We hypothesise that this is because the Netherlands is a very wet country with a huge number of very small water bodies, particularly in the west, meaning that there are few areas which are far from water (Het Waterschapshuis, 2017). The high abundances of mallards in the west also fits with the results of Barker et al. (2014), who found that mallards prefer areas with high water body density and open grassland. The preference of house sparrows for urban areas has been well documented. They nest in buildings and urban green spaces provide good food availability (Bernat-Ponce et al., 2018; Murgui, 2009; Robinson et al., 2005; Šálek et al., 2015). Ramírez-Cruz & Ortega-Álvarez (2021) also note that house sparrows likely avoid natural areas with high vegetation due to limited food resources, difficulty spotting predators and competition with other species. Their preference for the eastern parts of the country over the west could be attributed to the sandier soil in the east, since house sparrows like to take sand baths (Hauser, 1957; Pandian, 2023).

Blackbird abundance is expected to stay roughly the same in the future, while the abundance of mallards and house sparrows is expected to increase. Looking at the partial dependence plots (see supplementary materials), the largest differences between the three species are in the vegetative cover, particularly at the smaller buffer sizes. It is possible that blackbirds are responding differently to future changes in grass and tree cover than mallards and house sparrows and this is why we see the different changes in abundance. Since the vegetative cover variables were found to be particularly important, and all three species respond similarly to changes in climate and land use, this seems to be the most plausible explanation. There are only small differences in bird abundance between the four scenarios. This is possibly because we only look thirty years into the future, which means they have not had so much chance to diverge. Also, all the scenarios have similar trends: warmer temperatures, less agriculture and increasing urbanisation, but to different extents. It therefore makes sense that we see similar patterns in all the scenarios. This is helpful for controlling disease risk, since we have a good idea how bird populations will change, regardless of the scenario, and can plan accordingly.

In terms of what this means for disease risk, there are many factors to consider. All the three species studied had a positive relationship between abundance and urban areas and the Netherlands is expected to become more urban in future. Research on avian disease in urban areas has produced mixed results. Several studies have shown reduced pathogen prevalence in urban blackbirds when compared with their rural counterparts (Evans et al., 2009; Geue & Partecke, 2008). On the other hand, other studies have found that while avian disease risk is reduced in cities for some diseases, it increases for others (Martin & Boruta, 2014), for example for flaviviruses such as WNV, which can be transmitted by all three species considered here. Figuerola et al. (2024) propose that urbanisation provides opportunities for different rather than reduced pathogen transmission, as well as facilitating new interactions between vectors, hosts and parasites. The abundance of mallards and house sparrows is predicted to increase, providing additional potential disease hosts. It therefore seems prudent to regularly monitor bird populations in urban areas for diseases, particularly for flaviviruses and other pathogens which are known to benefit from an urban environment.

Avian abundance has been found to be positively related to disease prevalence for several pathogens, though there are exceptions and it should not be assumed that this will always be the case (Ellis et al., 2017; Santiago-Alarcon et al., 2016; States et al., 2009; Zhang et al., 2014). The increase in mallard abundance could have a particularly large effect, since they travel relatively large distances both nationally due to breeding dispersion (up to 19km) and internationally through migration, leading to a greater risk of diseases spreading to new areas (Bartel et al., 2018; van Turnhout, 2011). The highest increases in abundance were in areas which currently have low abundance, suggesting an ‘evening-out’ effect. This could potentially make it harder to predict areas which are at particularly high risk of disease outbreaks and necessitate more widespread surveillance than would otherwise be the case. There are also ongoing changes in bird migration patterns due to climate change, which we have not considered in this study. Both migration times and destinations are changing, potentially exposing birds to pathogens they would not otherwise have been exposed to (Hall et al., 2016; Pautasso, 2012). These changes create additional challenges for surveillance programmes, but also make such programmes all the more important. Increased host abundance, combined with higher uncertainty on potential outbreak locations and greater risk of new disease introductions, make widespread avian disease surveillance vital if new outbreaks are to be detected in a timely manner.

## Conclusions

Blackbirds, mallards and house sparrows are all likely to be especially relevant for disease transmission in Northwestern Europe and are capable of transmitting a wide range of pathogens. Their abundances will likely increase in future, especially for mallards and house sparrows. Unfortunately it is not possible to extrapolate these results to other bird species, since they will respond differently to future climate and land cover changes. However, it is interesting to note that while many bird species are expected to decline in future (Huntley et al., 2008; Rigal et al., 2023; Soultan et al., 2022), these three species which are potentially key for disease transmission will thrive. Quantifying the consequences of these abundance changes is complicated as there are many factors to consider, however it is likely that increased pathogen reservoirs will increase disease risk and that changes in distribution may affect local outbreak risk. The detailed future abundance maps created in this study will be a useful tool for disease modellers to estimate future risk.

## Data availability

All model inputs (with the exception of the bird data), model outputs, code and supplementary information can be accessed via the Dryad repository: https://doi.org/10.5061/dryad.r2280gbmc. The bird data is not publicly available but can be requested from the Dutch Centre for Field Ornithology (Sovon: sovon.nl) or Vigie-Nature (https://www.vigienature.fr/fr/suivi-temporel-des-oiseaux-communs-stoc). A summary of the bird data is available via the above Dryad link.

## Acknowledgements

We would like to thank Gerard van der Schrier for his advice on meteorological data sources; Vigie-Nature and the LPO (Ligue pour la Protection des Oiseaux) for developing and managing the French bird data collection program and all the volunteers who made the project possible; the thousands of volunteers of Sovon who go out to collect Dutch bird data; the E-OBS dataset from the EU-FP6 project UERRA (https://www.uerra.eu) and the Copernicus Climate Change Service, and the data providers in the ECA&D project (https://www.ecad.eu). This publication has been prepared using European Union’s Copernicus Land Monitoring Service information. This publication is part of the project ‘Preparing for vector-borne virus outbreaks in a changing world: a One Health Approach’ (NWA.1160.1S.210) which is (partly) financed by the Dutch Research Council (NWO).

## Notes

https://doi.org/10.5061/dryad.r2280gbmc

### Competing Interest Statement

The authors have declared no competing interest.

